# Molecular and Morphological Evidence Reveals *Cryptonema producta* in Shui Hau, Hong Kong, Previously Misidentified as *Anomalocardia flexuosa*

**DOI:** 10.64898/2026.05.14.725093

**Authors:** Lam Horace, Shen Lin, Ziyue Xu, Cynthia Sin Ting Yau, Longjun Wu

## Abstract

For over four decades, the bivalve *Anomalocardia flexuosa* has been recorded in Hong Kong coastal waters. However, the known native distribution of this heavily exploited commercial species is restricted to the Atlantic coast of South America, raising questions about the biogeographical validity of the Hong Kong populations. By employing an integrative taxonomic approach combining morphological re-evaluations and molecular phylogenetic analysis of the COI gene, we confirm that the species in Shui Hau, Hong Kong, China, has been historically misidentified. The population belongs to *Cryptonema producta* (syn. *Anomalocardia producta*).

## Introduction

The bivalve *Anomalocardia flexuosa* has been recorded in Hong Kong coastal waters for over 40 years, first appearing in Morton and Morton’s (1983) seminal field guide on the intertidal ecology of Hong Kong, and in more recent ecological surveys in Shui Hau (So et al., 2021; Lam et al., 2023), a relatively pristine sandflat in southwestern Hong Kong that is popular with recreational clam diggers. However, the presence of this species in Asia presents a significant geographical puzzle. The original description by Linnaeus (1767) was based on a specimen from the West Indies, and recent studies restrict its native range to the Atlantic coast of South America, extending from the Antilles to southern Brazil (de Oliveira Lima Gomes et al., 2019; Marinho et al., 2020; Bruzaca et al., 2022; Lopes et al., 2022). In Brazil, it is the most consumed and commercially harvested marine bivalve along the coast (Marinho et al., 2020). Interestingly, while it is included in the China Mollusca list (Chen et al., 2011), there are no records of it being commercially harvested in China. Furthermore, it is highly unusual for a coastal marine species to naturally exhibit such a vast, trans-oceanic gap in its distribution.

Compounding this geographical inconsistency are clear physical differences in the shell appearance. Authentic Brazilian *A. flexuosa* are generally equilateral in shape, sometimes exhibiting a yellowish colour to the external shell surface (Figure 1), with a series of robust concentric growth rings but no radial ribs. In contrast, the specimens collected from Shui Hau completely lack this yellow coloration and feature a distinctly extended posterior tip, together with fine concentric rings intersected by very fine radial ribs. These major discrepancies in geography, commercial utility, and shell morphology prompt the question of whether the specimens in Hong Kong have been misidentified for decades.

**Figure 1A:**
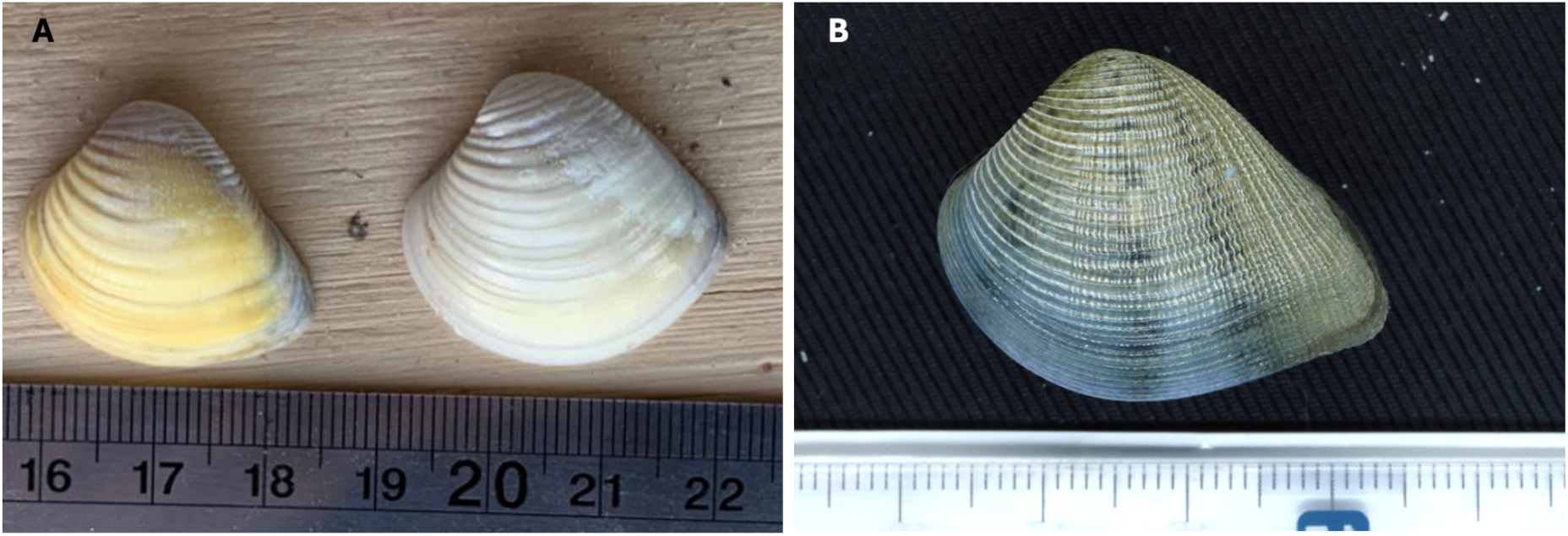
*A. flexuosa* from Brazil (Photo 254478169, jlcartes, 2023, iNaturalist). Figure 1B: “*A. flexuosa*” collected in Shui Hau, Hong Kong

## Methods

### Sample collection

Field surveys were conducted in August 2024 and March 2025 to collect live specimens of putative *A. flexuosa* from the intertidal zone of the sandy bay at Shui Hau (22°21’ 95.89’’ N,113°92’04.09’’E), Lantau Island, Hong Kong, China. Additional site visits were carried out at locations where the species had been historically recorded, specifically Starfish Bay in the northeast and Tai Tam in the south of Hong Kong. However, neither live individuals nor empty shells were found at these secondary sites. The live specimens collected from Shui Hau were transported to the laboratory and stored in a −20 °C freezer for subsequent DNA extraction, while the empty shells were cleaned and retained for morphological analysis.

### Morphological analysis DNA Extraction and Amplification

Mantle tissue samples (<30 mg) were excised from the clams for genomic DNA extraction using the TIANamp Marine Animals DNA Extraction Kit (Tiangen, Shanghai, China). Following extraction, the DNA was preserved in TE buffer and diluted to 100 μg/mL for polymerase chain reaction (PCR). The mitochondrial cytochrome c oxidase subunit I (COI) was selected as the target gene barcode (Chen et al., 2009; Zhang et al., 2012; Zheng et al., 2019; Feng et al., 2020; Chan et al., 2021; Hsiao & Chuang, 2023). Amplification was performed using the universal primers LCO-1490 (5′ GGTCAACAAATCATAAAGATATTGG-3′) and HCO-2198 (5′-TAAACTTCAGGGTGACCAAAAAATCA-3′). The 25 μl PCR mixture consisted of 2 μl of template DNA, 1 μl of each primer (10 μmol/L), 12.5 μl of Ex Taq polymerase version 2.0 (TaKaRa), and 8.5 μl of deionized water. The thermocycling profile included an initial denaturation at 94 °C for 3 min, followed by 35 cycles of denaturation at 94 °C for 40 s, annealing at 50 °C for 30 s, and extension at 72 °C for 45 s, with a final extension step at 72 °C for 5 min. PCR products were subsequently held at 15 °C. Success of the amplification was verified via electrophoresis on a 1% agarose gel. Target amplicons were excised and purified using the TaKaRa MiniBEST Agarose Gel DNA Extraction Kit Ver. 4.0 prior to sequencing by Sangon Biotech (Shanghai) Co., Ltd.

### Phylogenetic Analysis

The resulting COI sequences were compared against existing records in GenBank using the NCBI BLAST tool (Feng et al., 2020). Sequences for targeted and outgroup species were downloaded for phylogenetic reconstruction (Appendix 1). Maximum Likelihood (ML) phylogenetic analyses were implemented with IQ-TREE 2.4.0 (Minh et al., 2020). Firstly, to select an evolutionary model backbone (exchangeability matrix and Rate Heterogeneity Across Sites (RHAS) model (Yang, 1994)) and reconstruct an initial tree for subsequent analysis, an initialization step was performed using ModelFinder (Kalyaanamoorthy et al., 2017) implemented in IQ-TREE. The selected best model is HKY+F+I+G4. Then, the selected model backbone was used to reconstruct the phylogenetic tree. Ultrafast bootstrap test (UFBoot) with 1000 replicates was conducted to calculate bootstrap values and assess clade support.

## Results

### Morphological Analysis

A brief description of the shell morphology from putative *Anomalocardia flexuosa* in Shui Hau is provided as follows: the shell is relatively small, typically ranging from 3 to 4 cm in length. It is solid, inequilateral, and generally sub-triangular in shape, characterized by a fairly straight dorsal margin and a posterior margin that is considerably longer than the anterior. Notably, the posterior margin is strongly produced into an acute tip, whereas the anterior margin is broadly rounded. The ventral margin is entirely smooth, lacking any crenulations, and features a slight indentation immediately prior to the posterior tip. Externally, the shell surface is sculptured, possessing fine concentric growth rings intersected by very fine radial ribs (or riblets) across the posterior half of the shell, being most densely packed on the posterior acute portion of the shell. Shell coloration varies widely from a creamy white to a bluish-gray, occasionally exhibiting distinct blue pigmentation concentrated on the ventral edge. A prominent diagnostic feature is the presence of three distinct dark blue or gray stripes radiating from the umbo to the ventral margin. A conspicuous oval lunule is present on the anterior margin.

The original diagnosis of *A. flexuosa* by Linnaeus (1767) lacks sufficient detail for definitive species identification. Consequently, a comprehensive study by Narchi (1972), published under the unaccepted junior subjective synonym *A. brasiliana*, was utilized as the primary morphological reference. It is worth noting that Narchi’s (1972) account contains internal contradictions; the text describes an equilateral shell with a “deep pallial sinus,” whereas the accompanying illustrations depict an inequilateral shell with a very shallow sinus. Regardless of these discrepancies, neither the written description nor the illustrations match the specimens collected in the present study. As shown in Figure 2, the putative *A. flexuosa* from Shui Hau exhibit a distinct protruding posterior side forming an acute angle, a very deep pallial sinus, and a complete lack of crenulations on the free margin. This major morphological divergence strongly supports the hypothesis that the Hong Kong populations have been misidentified.

**Figure 2.**
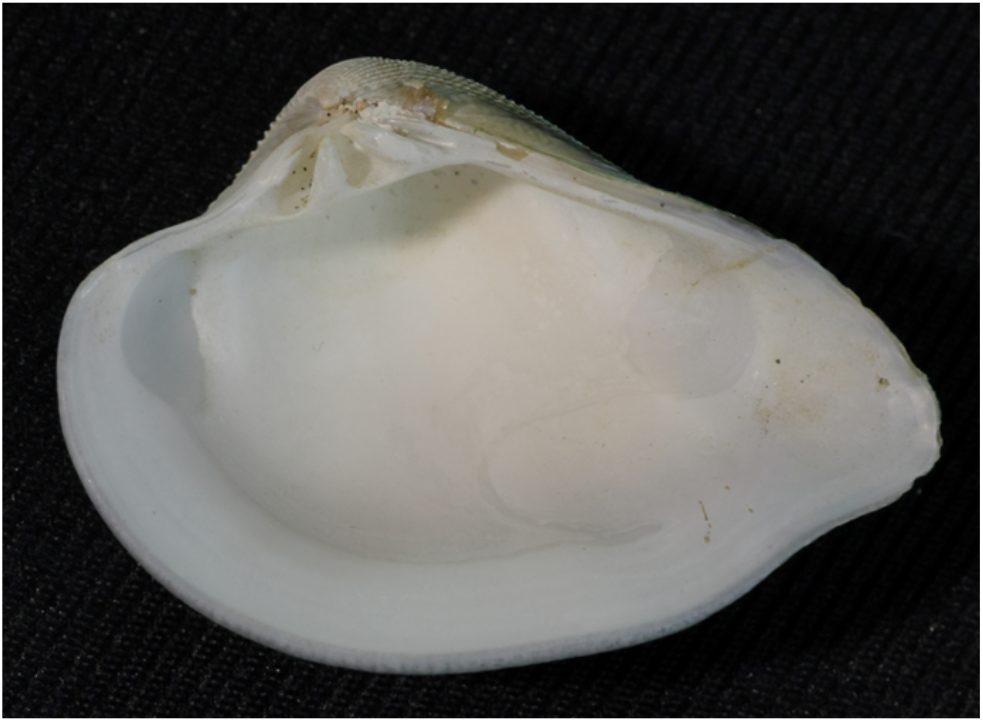
Internal view of “*A. flexuosa*” collected from Shui Hau, showing a very deep pallial sinus and no crenulations on the free margin.

Based on regional distributions, a likely candidate for the true identity of the Shui Hau specimens is *Cryptonema producta*. Originally described as *Venus impressa*, this species is native to Taiwan and the South China coast from Xiamen to Hainan (Xu, 1997). The original morphological description provided by Anton (1837) aligns closely with the Shui Hau samples on several key diagnostic features. Specifically, the protruding posterior margin and the distinctive pattern of “three olivaceous radial rays” on the outer surface are clearly present in the Shui Hau specimens. While the Shui Hau samples do exhibit two prominent cardinal hinge teeth consistent with Anton’s description, it is worth noting that the presence of an additional small anterior lateral tooth introduces a minor diagnostic discrepancy, highlighting the need for molecular validation.

### Molecular Phylogenetics

The partial COI gene from three putative *A. flexuosa* specimens collected at Shui Hau (samples 1, A10, and N1) was successfully sequenced. A BLAST search of these sequences yielded a 99% identity match with *Cryptonema producta* (syn. *Anomalocardia producta*). To further resolve their taxonomic placement, a Maximum Likelihood phylogenetic tree was constructed (Figure 3). This analysis incorporated our three novel sequences alongside reference COI sequences downloaded from GenBank for *C. producta*, true *A. flexuosa* (syn. *A. brasiliana*), and *Anomalocardia squamosa*. Node support was evaluated using bootstrap method. In the resulting phylogeny, all three putative *A. flexuosa* Shui Hau samples clustered tightly within the *C. producta* clade, confirming that they belong to the same species. Conversely, the reference sequences for authentic *A. flexuosa* formed a distinctly separate clade.Crucially, the clades representing *C. producta*, true *A. flexuosa* (syn. *A. brasiliana*), and *A. squamosa* were supported by high bootstrap values of 95, 100, and 99, respectively, providing robust statistical confidence that these represent distinct species. These molecular findings corroborate the morphological assessment, offering strong evidence that the Shui Hau population in Hong Kong is in fact, *C. producta*, and has been erroneously recorded as *A. flexuosa* in the local literature for decades.

**Figure 3:**
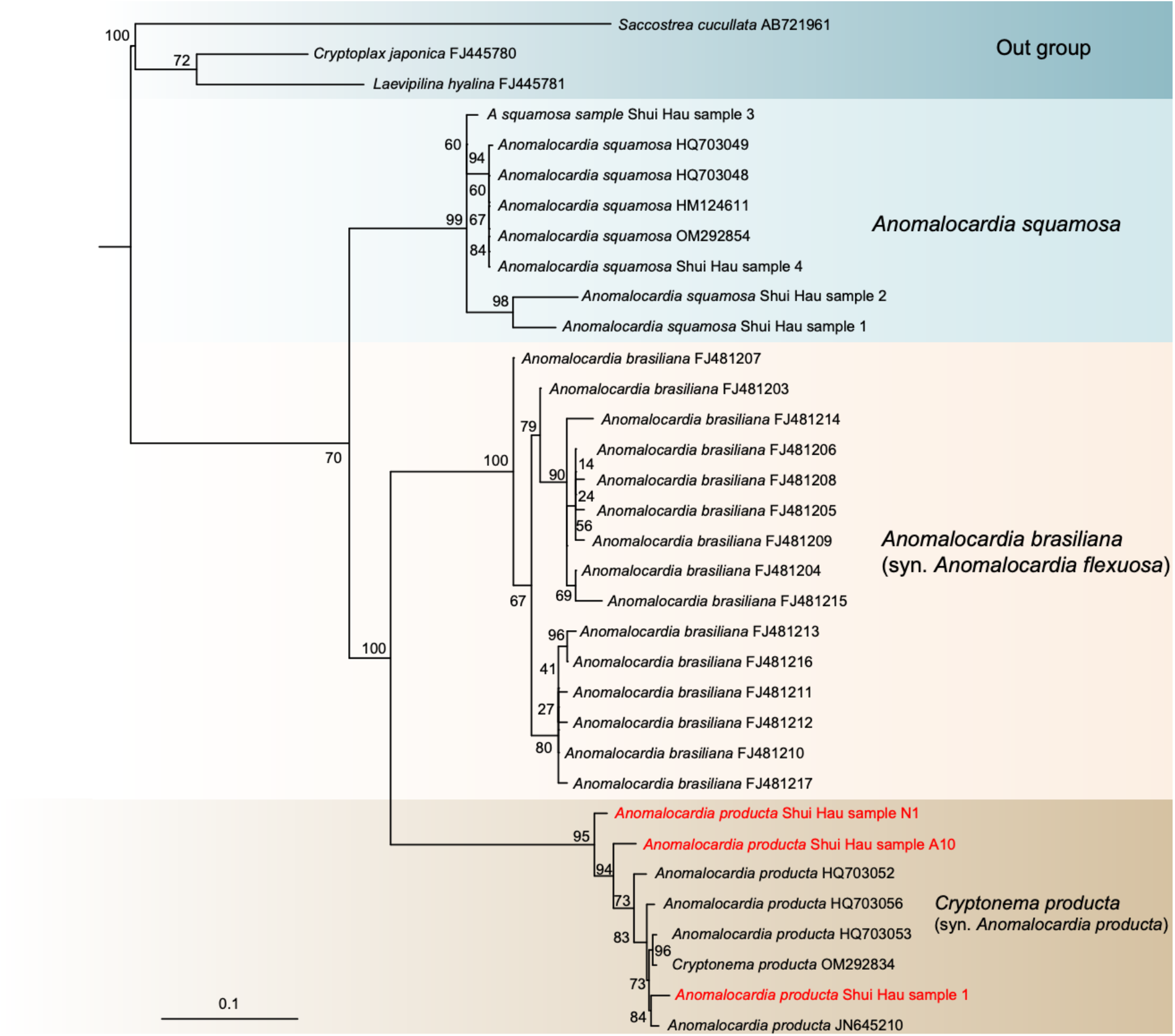
Phylogenetic tree of COI gene constructed using the maximum likelihood method. *Laevipilina hyaline, Cryptoplax japonica* and *Saccostrea cucullata* were used as outgroups of the tree.

## Conclusion

Based on the congruency of both morphological re-evaluations and molecular phylogenetic analyses, this study provides conclusive evidence that the bivalve historically recorded as *Anomalocardia flexuosa* in Shui Hau, Hong Kong, has been subject to a long-standing taxonomic misidentification. We verify that the true identity of this local population is *Cryptonema producta* (Kuroda et Habe, 1951).

## Future directions

Future research should prioritize expanding the sample size of *C. producta* for molecular analysis to substantiate these initial taxonomic findings. Additionally, a comprehensive ecological survey of this species throughout Hong Kong is urgently needed. Because neither live individuals nor empty shells were found at historically recorded locations such as Tai Tam and Starfish Bay during the present study, it is highly probable that local populations have declined or faced localized extirpation. Subsequent surveys should encompass other sandflat and mudflat habitats, such as Mui Wo and Hoi Ha Wan, where the putative *Anomalocardia flexuosa* was previously documented. Updating the contemporary distribution of this species in Hong Kong is highly critical, particularly for clarifying the impacts of recreational clam harvesting pressure on local population densities (So et al., 2021).

Furthermore, sampling efforts must be expanded across the broader geographical range of the species. Specimens should be collected from Guangxi and Hainan, where regional researchers have documented similar occurrences (Liu et al., 2017; Tinghe et al., 2019; Xu et al., 2024). Analyzing regional material is essential to confirm whether the *C. producta* populations in Hong Kong are conspecific with those in surrounding regions, and to determine if similar historical misidentifications have occurred elsewhere. Ultimately, a broad-scale phylogeographic assessment will confirm whether a single, widespread species of *C. producta* ranges across Southern China and Southeast Asia, or if cryptic diversity exists within the region.

## Appendix 1: Table of Gene sequences used in this study

|  |  |  |
| --- | --- | --- |
| *Anomalocardia producta* Shui Hau 1 | / | This study |
| *Anomalocardia producta* Shui Hau A10 | / | This study |
| *Anomalocardia producta* Shui Hau N1 | / | This study |
| *Anomalocardia producta* | JN645210 | GenBank |
| *Cryptonema producta* | OM292834 | GenBank |
| *Anomalocardia producta* | HQ703053 | GenBank |
| *Anomalocardia producta* | HQ703056 | GenBank |
| *Anomalocardia producta* | HQ703052 | GenBank |
| *Anoalocardia squamosa* Shui Hau 1 | / | This study |
| *Anoalocardia squamosa* Shui Hau 2 | / | This study |
| *Anoalocardia squamosa* Shui Hau 3 | / | This study |
| *Anoalocardia squamosa* Shui Hau 4 | / | This study |
| *Anomalocardia squamosa* | MN608368 | GenBank |
| *Anomalocardia squamosa* | MN608367 | GenBank |
| *Anomalocardia squamosa* | OM292854 | GenBank |
| *Anomalocardia squamosa* | HM124611 | GenBank |
| *Anomalocardia squamosa* | HQ703048 | GenBank |
| *Anomalocardia squamosa* | HQ703049 | GenBank |
| *Anomalocardia brasiliiana* | DQ458471 | GenBank |
| *Anomalocardia brasiliiana* | FJ481217 | GenBank |
| *Anomalocardia brasiliiana* | FJ481210 | GenBank |
| *Anomalocardia brasiliiana* | FJ481212 | GenBank |
| *Anomalocardia brasiliiana* | FJ481211 | GenBank |
| *Anomalocardia brasiliiana* | FJ481216 | GenBank |
| *Anomalocardia brasiliiana* | FJ481213 | GenBank |
| *Anomalocardia brasiliiana* | FJ481215 | GenBank |
| *Anomalocardia brasiliiana* | FJ481204 | GenBank |
| *Anomalocardia brasiliiana* | FJ481209 | GenBank |
| *Anomalocardia brasiliiana* | FJ481206 | GenBank |
| *Anomalocardia brasiliiana* | FJ481208 | GenBank |
| *Anomalocardia brasiliiana* | FJ481205 | GenBank |
| *Anomalocardia brasiliiana* | FJ481207 | GenBank |
| *Anomalocardia brasiliiana* | FJ481203 | GenBank |
| *Anomalocardia brasiliiana* | FJ481214 | GenBank |
| *Laevipilina hyalina* | FJ445781 | GenBank |
| *Cryptoplax japonica* | FJ445780 | GenBank |
| *Saccostrea cucullata* | AB721961 | GenBank |

## Appendix 2: Sequences used in this study

### >Cryptonema_producta_Shui_Hau_1

TCTCCTTTTTTCATTTGAGCTGGGCTTATAGGAACGGCTTTTAGTGTTATTATTCGA ATGGAGTTAGCTATACCAGGAAAGATGCTCGATGATGGACAGTTATATAATTTAGTT GTTACTGCTCATGGTTTAGTTATAATTTTCTTTTTAGTAATACCTATGTTAATTGGTG GGTTTGGAAATTGGTTAATTCCTCTTATACTAAAAATCCCTGATATAGCTTTTCCAC GAATAAATAACTTAAGATTCTGATTGCTACCAGTTTCTATATTACTTTTGTTAGGGTC CTCTTATGTAGATGGAGGTGCGGGAACGGGTTGAACAGTATATCCACCTTTATCTTC TATTTTATTTCACTCAGGTTGTGCTGTTGATTATGTGATTTTTTCACTGCACGTGGGT GGTGTATCATCAATTTTGGCTTCTATTAATTTTGCAATTACTACGGCATTAATACGAA CTAGAGTAATAATTATTCTGCGTAGAAGAATGTTGGCTTGGTGTTTAGGGATTACTA GGTTTCTTTTAATTGTAGCTATACCAGTATTGGCAGGAGGTTTAACTATGCTTTTAA CTGATCGTCATTTTAATACTACATTTTTTGACCCAATAGGAATGGGGGATCCAATTC TATTTATTCACTTATTTTGATTTTTTGGTCACCCTGGAAGTTTAAA

### >Cryptonema_producta_Shui_Hau_A10

GATCTTTTATTTTTTCATTTGAGCTGGACTCATAGGAACGGCTTTTAGTGTTATTATTCGAATGGAGTTAGCTATACCAGGAAAGATGCTCGATGATGGACAGTTATATAATTTAGTTGTTACTGCTCATGGTTTAGTTATAATTTTCTTTTTAGTAATACCTATGTTAATTGGTGGGTTTGGAAATTGATTAATTCCCCTTATACTAAAAATTCCTGATATAGCTTTTCCACGAATAAATAACTTAAGATTCTGATTGCTTCCAGTTTCTATATTACTCTTGTTAGGGTCTTCTTATGTAGATGGGGGTGCGGGAACGGGTTGAACAGTATATCCACCTTTATCTTCTATTTTATTTCACTCAGGTTGTGCTGTTGATTATGTGATTTTTTCACTGCACGTGGGTGGTGTATCATCAATTTTGGCTTCTATTAATTTTGCAATTACTACGGCATTAATACGAACCAGAGTAATAATTATTCTGCGTAGAAGAATGTTGGCTTGGTGTTTAGGGGTTACTAGGTTTCTTTTAATTGTAGCTATACCAGTATTGGCAGGAGGTTTAACTATGCTTTTAACTGATCGTCATTTTAATACTACATTTTTTGACCCAATAGGAATGGGGGACCCAATTCTATTTATTCACTTATTTTGATTTTTTGGTCACCCTGGAAGTTTAAA

### >Cryptonema_producta_Shui_Hau_N1

TGGTCAACCCAAACATAAAGATATTGGTACATTATATTTTATTTTTTCGATTTGAGCTGGACTTATAGGAACGGCTTTTAGTGTTATTATTCGAATGGAATTAGCTATACCAGGAAAGATGCTCGATGATGGACAGTTATATAATTTAGTTGTTACTGCTCATGGTTTAGTTATAATTTTCTTTTTAGTAATACCTATGTTAATTGGTGGGTTTGGAAATTGATTAATTCCTCTTATACTAAAAATTCCTGATATAGCTTTTCCACGAATAAATAACTTAAGATTCTGATTGCTACCAGTTTCTATATTACTCTTGTTAGGATCCTCTTATGTAGATGGGGGTGCGGGAACGGGTTGAACAGTATATCCACCTTTGTCTTCTATTTTATTTCACTCAGGTTGTGCTGTTGATTATGTGATTTTTTCACTGCACGTGGGTGGTGTATCATCAATTTTGGCT TCTATTAATTTTGCAATTACTACGGCATTAATACGAACCAGAGTAATAATTATTCTGCGTAGAAGAATGTTGGCTTGGTGTTTAGGGGTTACTAGGTTTCTTTTAATTGTAGCTATACCAGTATTGGCAGGAGGTTTAACTATGCTTTTAACTGATCGTCATTTTAATACTACATTTTTTGACCCAATAGGAATGGGGGACCCAATCTATTTATTCACCATT

### *>Anoalocardia squamosa* Shui Hau 1

TGTTTCCAAACACAAACATAAAGATATTGGAACTTTATACTTTATTTTTTCAATTTGATCTGGTTTGATGGGAACTGCTTTTAGTGTAATTATGCGAATAGAGTTGGCTATGCCTGGAAAGATGTTGGATGATGGTCAATTATATAATTTAATTGTTACGGCTCATGGGTTGGTAATAATTTTTTTTCTTGTTATACCAATAATAATTGGGGGTTTTGGTAACTGGTTGGTACCTTTGATGTTAACTATACCTGATATGGCTTTCCCTCGAATAAATAATTTAAGCTTTTGATTATTACCTGTTTCGATAGCTTTATTATTAAGTTCTAGTTATGTAGATGGGGGTGCCGGAACTGGTTGAACTATTTATCCTCCTTTATCTAGAAATATGTCTCACTCTGGTCCTGCTATAGATTATGTTATTTTTTCCTTACACTTGGGGGGTGCTTCATCAATTATGGCTTCTATTAATTTCGTTTGTACTTGTTTCTTAATGCGTCCAGGAGTGATAACTTTGTTGCGCACCTCGATGTTTTGTTGATGTGTTAGGGTCACTGGGTTTTTACTGGTTGTTGCAATACCTGTTCTAGCTGGTGCATTAACTATGTTGTTGACCGTATCGAAATTTTAACACGTGTCTATACTTTACAGACCTGCAGTAGGAGGCACGACGCCATCCTCTTTTCCA

### *>Anoalocardia squamosa* Shui Hau 2

TGGTCCAAACCAAAACATAAAGATATTGGAACTTTATACTTTATTTTTTCAATTTGATCTGGTTTGATGGGAACTGCTTTTAGTGTAATTATGCGAATAGAGTTGGCTATGCCTGGAAAGATGTTGGATGATGGTCAATTATATAATTTAATTGTTACGGCTCATGGGTTGGTAATAATTTTTTTTCTTGTTATACCAATAATAATTGGGGGTTTTGGTAACTGGTTGGTACCTTTGATGTTAACTATACCTGATATGGCTTTCCCTCGAATAAATAATTTAAGCTTTTGATTATTACCTGTTTCGATAGCTTTATTATTAAGTTCTAGTTATGTAGATGGGGGTGCCGGAACTGGTTGAACTATTTATCCTCCTTTATCTAGAAATATGTCTCACTCTGGTCCTGCTATAGATTATGTTATTTTTTCCTTACACTTGGGGGGTGCTTCATCAATTATGGCTTCTATTAATTTCGTTTGTACTTGTTTCTTAATGCGTCCAGGAGTGATAACTTTGTTGCGCACCTCGATGTTTTGTTGATGTGTTAGGGTCACTGGGTTTTTACTGGTTGTTGCAATACCTGTTCTAGCTGGTGCATTAACTATGTTGTTGACCGATCGAAATTTTAACATGTGTCTTTTTTCAGACCCATGTAGGAGGTACACGGCATACACTATGATCCAT

### *>Anoalocardia squamosa* Shui Hau 3

TGGTCAAACCAAACATAAAGATATTGGAACTTTATACTTTATTTTTTCAATTTGATCTGGTTTGATGGGAACTGCTTTTAGTGTAATTATGCGAATAGAGTTGGCTATGCCTGGAAAGATGTTGGATGATGGTCAATTATATAATTTAATTGTTACGGCTCATGGGTTGGTAATAATTTTTTTTCTTGTTATACCAATAATGATTGGGGGTTTTGGTAACTGGTTGGTACCTTTGATGTTAACTATACCTGATATGGCTTTCCCTCGAATAAATAATTTAAGCTTTTGATTATTACCTGTTTCGATAGCTTTATTATTAAGTTCTAGTTATGTAGATGGGGGTGCCGGAACTGGTTGAACTATTTATCCTCCTTTATCTAGAAATATGTCTCACTCTGGTCCTGCTATAGATTATGTTATTTTTTCCTTACACTTGGGGGGTGCTTCATCAATTATGGCTTCTATTAATTTCGTTTGTACTTGTTTCTTAATGCGTCCAGGAGTGATAACTTTGTTGCGCACCTCGATGTTTTGTTGATGTGTTAGGGTCACTGGGTTTTTACTGGTTGTTGCAATACCTGTTCTAGCTGGTGCATTAACTATGTTGTTGACCGATCGAAATTTTAACACTTCTTTTTTCGACCCCGTAGGAATAGGGGATCCAGTTA

### >*Anoalocardia squamosa* Shui Hau 4

TGTGTTACAACCAAAACATAAAGATATTGGAACTTTATACTTTATTTTTTCAATTTGATCTGGTTTGATGGGAACTGCTTTTAGTGTAATTATGCGAATAGAGTTGGCTATGCCTGGAAAGATGTTGGATGATGGTCAATTATATAATTTAATTGTTACGGCTCATGGGTTGGTAATAATTTTTTTTCTTGTTATACCAATAATAATTGGGGGTTTTGGTAACTGGTTGGTACCTTTGATGTTAACTATACCTGATATGGCTTTCCCTCGAATAAATAATTTAAGCTTTTGATTATTACCTGTTTCGATAGCTTTATTATTAAGTTCTAGTTATGTAGATGGGGGTGCCGGAACTGGTTGAACTATTTATCCTCCTTTATCTAGAAATATGTCTCACTCTGGTCCTGCTATAGATTATGTTATTTTTTCCTTACACTTGGGGGGTGCTTCATCAATTATGGCTTCTATTAATTTCGTTTGTACTTGTTTCTTAATGCGTCCAGGAGTGATAACTTTGTTGCGCACCTCGATGTTTTGTTGATGTGTTAGGGTCACTGGGTTTTTACTGGTTGTTGCAATACCTGTTCTAGCTGGTGCATTAACTATGTTGTTGACCGATCGAAATTTTAACA CTTCTTTTTTCGACCCCGTAGGAATAGGGGATCCTGTTTTATTATCAATT

